# A Shared Entropic Axis Spans States of Consciousness Across Pharmacological and Clinical Conditions

**DOI:** 10.64898/2026.06.12.731897

**Authors:** Dante Sebastián Galván Rial, Gabriel A. Della Bella, Lorina Naci, Stefano Delli Pizzi, Stefano L. Sensi, Tristán M. Osán, Daniel Aguilera, Christopher Timmermann, Robin Carhart-Harris, Emmanuel A. Stamatakis, Pablo Barttfeld

## Abstract

A general account of how diverse states of consciousness arise from brain activity requires a quantitative framework that generalises across them. The Entropic Brain Theory proposes that states of consciousness can be ordered along a single dimension defined by the entropy of spontaneous neural activity, but this prediction has not been tested across pharmacological and clinical perturbations within a common analytical pipeline. Here we quantify the temporal irregularity of time-resolved small-world topology using sample entropy, applying the same pipeline to pharmacological (psychedelics, modafinil, propofol anaesthesia) and clinical (schizophrenia) fMRI datasets. Propofol anaesthesia occupied the low-entropy end of the axis; psychedelic states and schizophrenia occupied the high end. The ordering tracks combined modulations of the level and content of consciousness, ranging from reduced awareness under anaesthesia to the heightened arousal and expanded experience of psychedelic states and the disorganised, dysregulated processing of schizophrenia. Crucially, this result was not reducible to fluctuations in mean functional connectivity, and was supported by convergent reorganisation of higher-order association cortex under psychedelics and anaesthesia, alongside a distributed loss of network specificity in schizophrenia. These findings provide cross-condition empirical support for an entropic continuum of brain states and identify the temporal diversity of large-scale network reconfiguration as a primary axis of conscious dynamics.

## 1 Introduction

A central problem in the neuroscience of consciousness is to identify a quantitative signature of brain dynamics that tracks the capacity for conscious experience. Two decades of theoretical and empirical work have linked conscious awareness to the large-scale organisation of brain activity, with successive proposals, from network efficiency and integration measures^1,2^ to dynamical signatures of psychedelic and meditative states^3,4^, converging on the idea that flexible reconfiguration of large-scale networks underlies conscious states. Measures of neural complexity have been operationalised differently across traditions, including algorithmic and perturbational measures from TMS-EEG^5,6^, graph-theoretical measures from fMRI^7^, signal diversity from M/EEG^8^, and entropy of dynamic functional connectivity^3,9^. Varley et al.^10^ showed that algorithmic, dynamical, and network-based indices all decrease under propofol, but this convergence has been demonstrated only within isolated paradigms, and the empirical case for a shared dynamical signature across states remains weak.

A second source of fragmentation is conceptual. Alterations of consciousness can affect either its level (the capacity for awareness) or its phenomenal content (the organisation of experience, including its coherence and self-referential framing^11^). These dimensions are dissociable in principle but coupled in practice, and perturbations engage them asymmetrically. Anaesthesia abolishes awareness itself, and with it any content, by collapsing level rather than by reorganising experience. Schizophrenia, by contrast, disrupts the organisation and self-referential framing of experience while the capacity for awareness is broadly retained^12^.

Psychedelics, in turn, modulate both dimensions: at moderate doses they elevate phenomenal richness alongside markers of autonomic arousal while basic reportability is largely retained, though higher doses can compromise reportability and awareness itself. A measure that generalises across these conditions must therefore track more than the level of awareness, and more than content alone: it must capture a property of brain dynamics that is altered whenever either dimension is perturbed.

The Entropic Brain Theory^13^ attempts this unification, positing that states of consciousness can be ordered along an entropic continuum. Subsequent empirical work, recently reviewed^14^, has been broadly consistent but variable in terms of entropy operationalisation (entropy of connectivity-degree distributions, signal diversity, local temporal structure) and conditions studied^3,7,8^. No study has tested this ordering across pharmacological and clinical conditions within a single pipeline, leaving the core prediction unresolved. Within this framework, small-world topology can be understood as the structural backbone that balances functional integration and segregation, allowing the brain to combine specialised local processing with efficient global communication^15^. The way this organisation fluctuates over time reflects how flexibly the brain moves between network configurations, and the temporal irregularity of small-worldness offers a tractable index of that flexibility: higher irregularity indicates a broader, more variable repertoire of network states, lower irregularity a more constrained one. Tracking how small-world organisation fluctuates over time thus links the abstract entropic account of consciousness to a measurable property of large-scale brain dynamics. We therefore propose that the property unifying these states is the temporal diversity of large-scale network reconfiguration, captured by a single index, the sample entropy of dynamic small-worldness. On this view, the heterogeneity of previously reported metrics reflects different windows onto one underlying dynamical constraint rather than a plurality of independent dimensions: states of consciousness are ordered along a single axis and perturbed in opposite directions from waking baseline, regardless of whether level, content, or both are affected.

We applied a single pipeline to five fMRI datasets—propofol anaesthesia, psychedelics (LSD, DMT), modafinil, schizophrenia, and matched controls— quantifying the temporal irregularity of large-scale network topology with sample entropy of time-resolved small-world dynamics. Brain states fell along an entropic gradient: anaesthesia at the low end, psychedelic and clinical conditions at the high end, healthy wakefulness in between. The ordering was independent of connectivity magnitude, and a network-level decomposition pointed to specific, condition-dependent network contributions, supporting the framework and positioning dynamic-network entropy as a robust index of conscious brain states.

## 2 Results

### 2.1 Condition-specific modulation of dynamic network topology

To ask whether dynamic-network entropy captures a property of brain dynamics shared across heterogeneous states of altered consciousness, we computed sample entropy on time-resolved trajectories of dynamic small-worldness (SE dSW) for each subject and condition (Fig. 1). SE dSW indexes how predictable the global network topology is from one moment to the next: high values reflect rich, non-repetitive trajectories through network space; low values, stereotyped and constrained dynamics. The measure is sensitive to the configuration of network connections rather than their strength; a complementary index based on the entropy of dynamic functional connectivity (SE dFC), introduced in a later section, tests whether topological effects can be reduced to fluctuations in connectivity magnitude.

**Figure 1.**
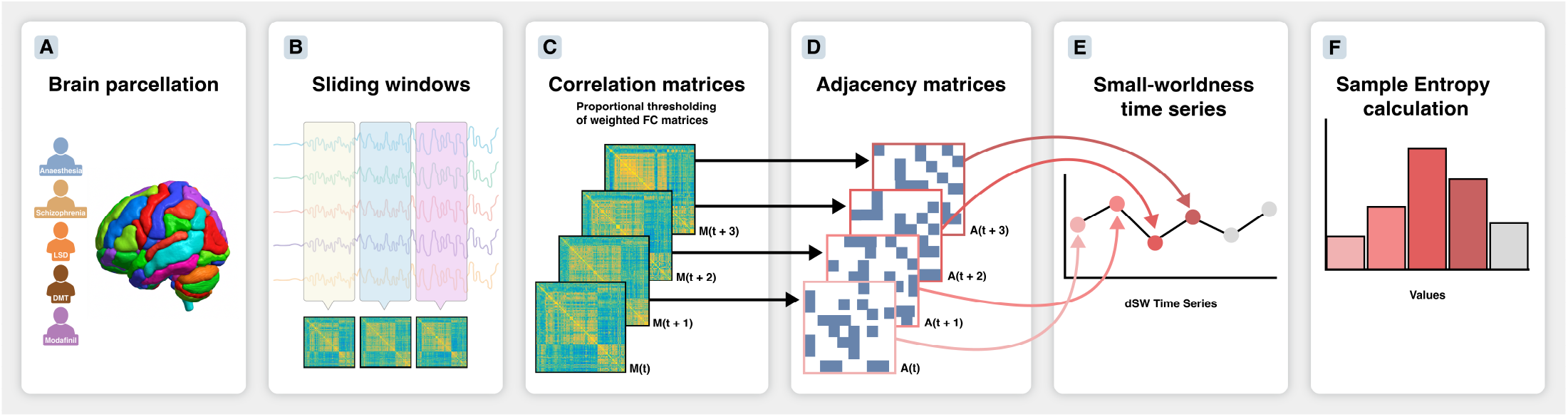
Analytical pipeline for dynamic-network entropy. For each of the five datasets, resting-state fMRI BOLD time series are extracted from parcellated regions of interest (A) and submitted to a sliding-window approach (B), producing a time-resolved functional connectivity matrix per window (C). Each matrix is proportionally thresholded, retaining edge weights, to yield a series of weighted adjacency matrices (D), from which dynamic small-worldness is computed at each time point, giving a small-worldness trajectory per participant and condition (E). Sample entropy (SE) is then applied to this trajectory to quantify its temporal irregularity (SE dSW; F). In parallel (not shown), SE is computed on the scalar dynamic functional connectivity (dFC) time series — the entropy of fluctuations in overall coupling strength — yielding SE dFC, a complementary measure that dissociates topological effects from changes in connectivity magnitude.

Propofol anaesthesia produced a graded suppression of network entropy (Fig. 2A). SE dSW decreased monotonically with depth of sedation: from awake baseline to light anaesthesia (*β* = -0.10, 95% CI [-0.19, -0.01], p = 0.036), and from awake baseline to deep anaesthesia (*β* = -0.14, 95% CI [-0.23, -0.04], p = 0.004). Upon recovery, entropy returned to baseline (*β* = -0.00, 95% CI [-0.10, 0.09], p = 0.95), indicating that the effect was reversible and tied to the active pharmacological state rather than to lasting effects of sedation. These results thus show pharmacological unconsciousness as a stereotyped, predictable mode of network reconfiguration.

**Figure 2.**
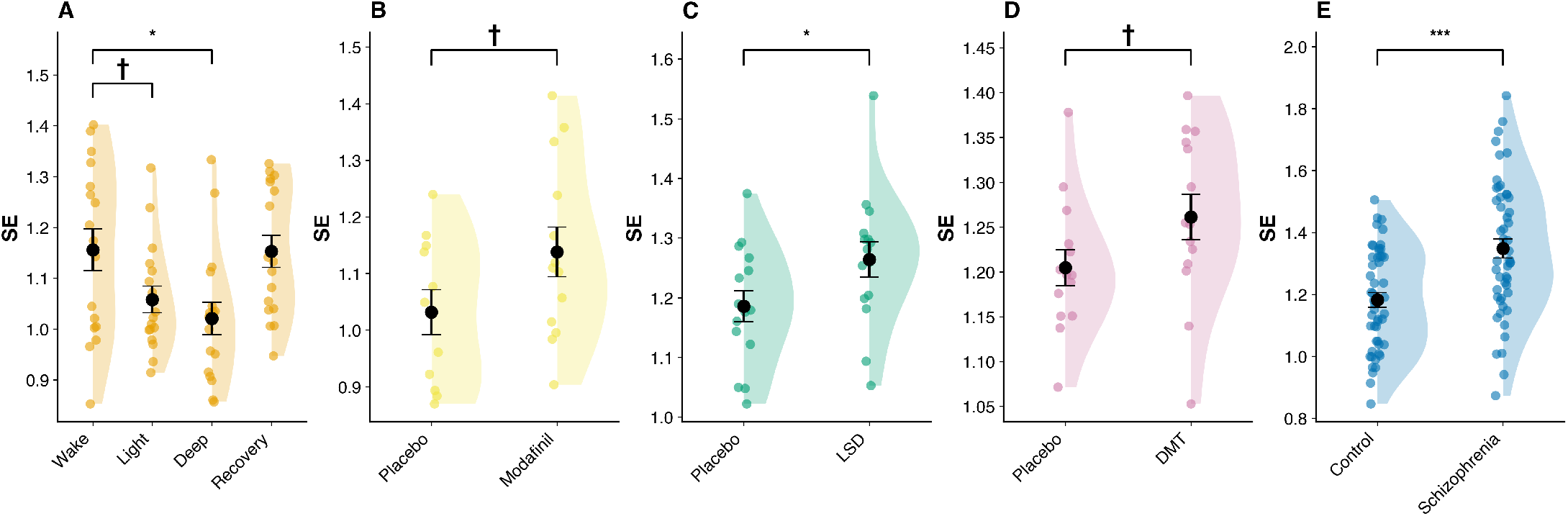
Dynamic small-worldness entropy (SE dSW) across individual datasets. Raincloud plots show the distribution of SE dSW for each independent study cohort. (A) Propofol anaesthesia: SE dSW decreased monotonically from awake baseline to light and deep anaesthesia, and returned to baseline upon recovery. (B) Modafinil: trend toward elevated SE dSW relative to placebo. (C) LSD: elevated SE dSW relative to placebo. (D) DMT: trend toward elevated SE dSW relative to placebo. (E) Schizophrenia: patients showed elevated SE dSW relative to matched healthy controls. Individual dots represent single subjects or sessions; central black circles denote sample means; error bars indicate s.e.m. †p < 0.10, *p < 0.05, **p < 0.01, ***p < 0.001.

A different pattern emerged for states that preserve awareness while altering its content or arousal. LSD elevated SE dSW relative to placebo (*β* = +0.08, 95% CI [0.01, 0.15], p = 0.028; Fig. 2C), and DMT showed a positive trend in the same direction (*β* = +0.06, 95% CI [-0.01, 0.12], p = 0.076; Fig. 2D). Modafinil produced a similar trend, placing it just above standard wakefulness (*β* = +0.11, 95% CI [-0.02, 0.23], p = 0.089; Fig. 2B). The largest elevation, however, was observed in schizophrenia: patients showed an increase in SE dSW relative to matched controls more than twice the magnitude of the psychedelic effects (*β* = +0.17, 95% CI [0.09, 0.24], p = 4.0 *×* 10^*−*5^; Fig. 2E). Together, these results show that elevations in topological entropy are not specific to any single class of perturbation: they appear under acute serotonergic agonism, under dopaminergic and noradrenergic modulation of arousal, and in a chronic clinical condition characterised by disorganised processing and altered self-organisation of experience.

### A macroscopic entropic gradient organises diverse states of consciousness

These condition-specific effects raised the question of whether the perturbations studied here could be embedded along a single ordered dimension of network entropy. To place the conditions on a common scale despite differences in acquisition site and baseline across studies, we standardised SE dSW within each study and expressed every perturbation as its standardised change relative to the corresponding within-study baseline (ΔSE dSW); we then fitted a linear mixed-effects model estimating each condition’s deviation from baseline wakefulness, with subject-level random intercepts to account for repeated measures (using raw values yields qualitatively identical results; Fig. S4). Ordering the perturbations by their mean ΔSE dSW reveals a continuous gradient (Fig. 3). Deep propofol anaesthesia anchors the low end, with the largest reduction (Coef = -0.80, 95% CI [-1.43, -0.17], p = 0.013); light anaesthesia shows a smaller, trend-level reduction (Coef = -0.58, 95% CI [-1.21, 0.05], p = 0.073); and recovery returns to baseline (Coef = -0.02, 95% CI [-0.65, 0.61], p = 0.96). Above baseline, LSD (Coef = +0.68, 95% CI [0.02, 1.34], p = 0.045), DMT (Coef = +0.83, 95% CI [0.15, 1.51], p = 0.017), and modafinil (Coef = +0.91, 95% CI [0.20, 1.61], p = 0.012) occupy progressively higher positions, and schizophrenia shows the largest elevation (Coef = +1.02, 95% CI [0.65, 1.38], p < 0.001).

**Figure 3.**
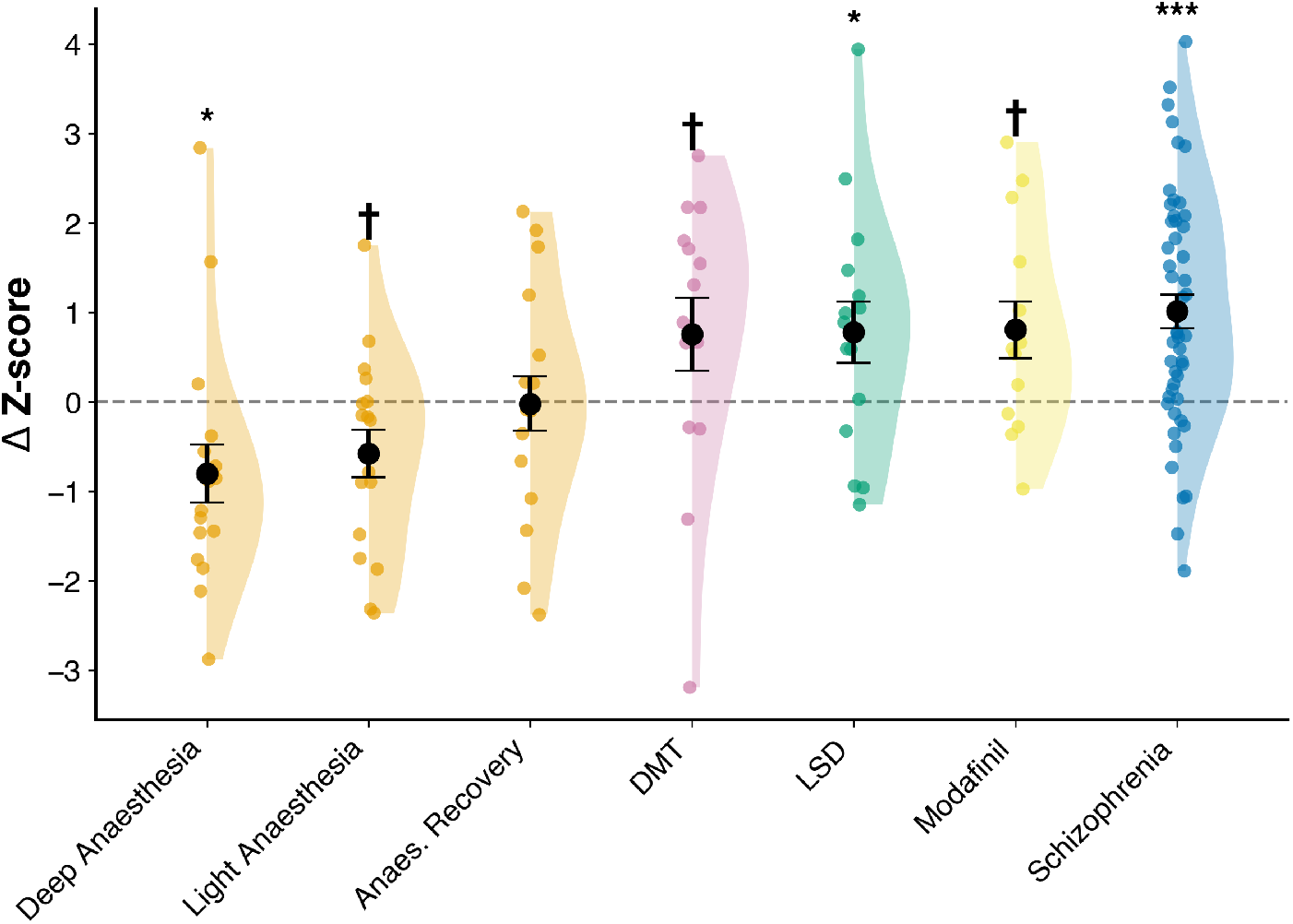
Continuous entropic gradient across perturbations of consciousness. For each perturbation, SE dSW was standardised within study, and its change relative to the within-study baseline is shown (ΔSE dSW; active minus control, or each sedation level minus awake baseline). Because SE dSW is standardised before differencing, values are expressed in standardised units (Δ Z-score, as labelled on the axis). Conditions are ordered by group mean from lowest to highest. The dashed line marks no change from baseline (Δ = 0). Deep and light propofol anaesthesia show the largest reductions, anaesthesia recovery returns to baseline, and DMT, LSD, modafinil and schizophrenia show progressive elevations, with schizophrenia the highest. Individual dots represent single subjects or sessions; black circles denote group means; error bars indicate s.e.m. †p < 0.10, *p < 0.05, **p < 0.01, ***p < 0.001.

Across the full set of perturbations, SE dSW thus orders states from suppressed to expanded network reconfiguration along a single continuous axis. This ordering was preserved when the entire pipeline was repeated with an alternative parcellation (Schaefer cortical with Melbourne subcortical; Fig. S2 for SE dSW, Fig. S3 for SE dFC).

### 2.3 Divergent effects on the entropy of global functional connectivity

The unified gradient is built on a measure of topological reconfiguration. But topology and the magnitude of connectivity are distinct properties of brain dynamics. The gradient observed above could therefore arise from genuine reorganisation of network configuration or, simply, from fluctuations in overall connectivity strength. In that case, it would reflect a byproduct of how strongly the brain is coupled rather than how it reconfigures. To test this, we computed sample entropy on the time series of mean dFC (SE dFC; Fig. 4). The two measures dissociated under psychedelics: both DMT (*β* = -0.06, 95% CI [-0.11, -0.00], p = 0.042, Fig. 4D) and LSD (*β* = -0.06, 95% CI [-0.09, -0.02], p = 0.001, Fig. 4C) reduced SE dFC relative to placebo, the opposite direction to their effect on SE dSW. Modafinil produced no detectable change in SE dFC (*β* = +0.00, p = 0.97, Fig. 4B). In the remaining conditions, the two measures moved together: SE dFC was suppressed by light and deep propofol anaesthesia (*β* = -0.06, p = 0.016 and *β* = -0.06, p = 0.012, Fig. 4A) and elevated in schizophrenia (*β* = +0.08, 95% CI [0.04, 0.12], p = 1.6 *×* 10^*−*4^, Fig. 4E), mirroring the SE dSW results in those datasets. Assembled into a gradient as before, the SE dFC values (Fig. S1) preserve the anaesthesia-low, schizophrenia-high structure seen for SE dSW, but the psychedelic conditions now fall at the low end of the axis —at or below the anaesthesia level— rather than at the high end they occupy for SE dSW, reflecting their opposite effects on the two measures: elevated topological entropy alongside reduced connectivity-magnitude entropy. The corresponding raw, un-differenced SE dFC values are reported in Fig. S5.

**Figure 4.**
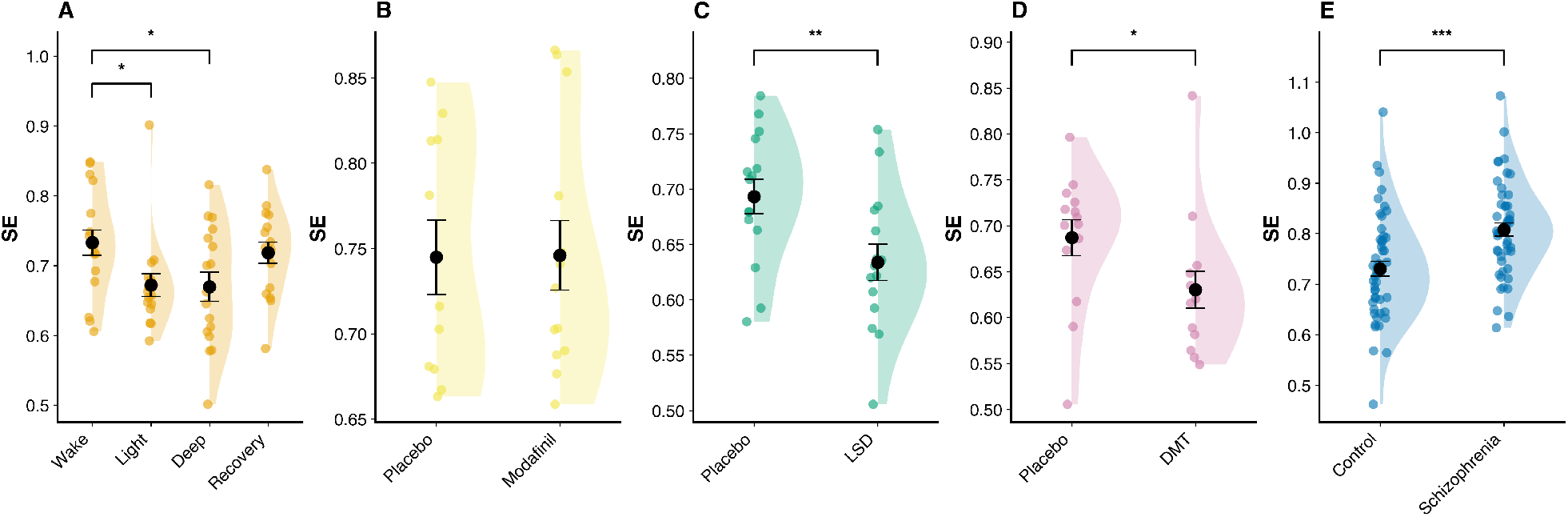
Dynamic functional connectivity entropy (SE dFC) across individual datasets. Raincloud plots show the distribution of SE dFC for each independent study cohort. (A) Propofol anaesthesia: SE dFC decreased monotonically with depth of sedation, and returned to baseline upon recovery. (B) Modafinil: no detectable change in SE dFC. (C) LSD: reduced SE dFC relative to placebo — the opposite direction to its effect on SE dSW (Fig. 2C). (D) DMT: reduced SE dFC relative to placebo — likewise opposite to its effect on SE dSW (Fig. 2D). (E) Schizophrenia: patients showed elevated SE dFC relative to matched healthy controls. Individual dots represent single subjects or sessions; central black circles denote sample means; error bars indicate s.e.m. *p < 0.05, **p < 0.01, ***p < 0.001.

The temporal diversity of network configurations (dSW) and the variability of connectivity magnitude (dFC) are therefore distinct properties of brain dynamics, with psychedelics modulating the former while suppressing the latter, and modafinil modulating the former without affecting the latter. Projected onto a two-dimensional state space defined by SE dSW and SE dFC (Fig. 5), psychedelic conditions occupy a region of the plane that no other perturbation reaches: high topological entropy combined with reduced connectivity entropy, distinct from the diagonal trajectory of propofol (low-low) and schizophrenia (high-high). The entropic gradient identified above thus describes one principal axis of a multidimensional dynamical space; conditions that share a position along that axis can still differ in how their topological dynamics relate to their connectivity dynamics. The same pattern of condition positions in the SE dSW *×* SE dFC plane was obtained using an alternative parcellation (Schaefer cortical with Melbourne subcortical; Fig. S6), confirming that the dissociation does not depend on parcellation choice.

**Figure 5.**
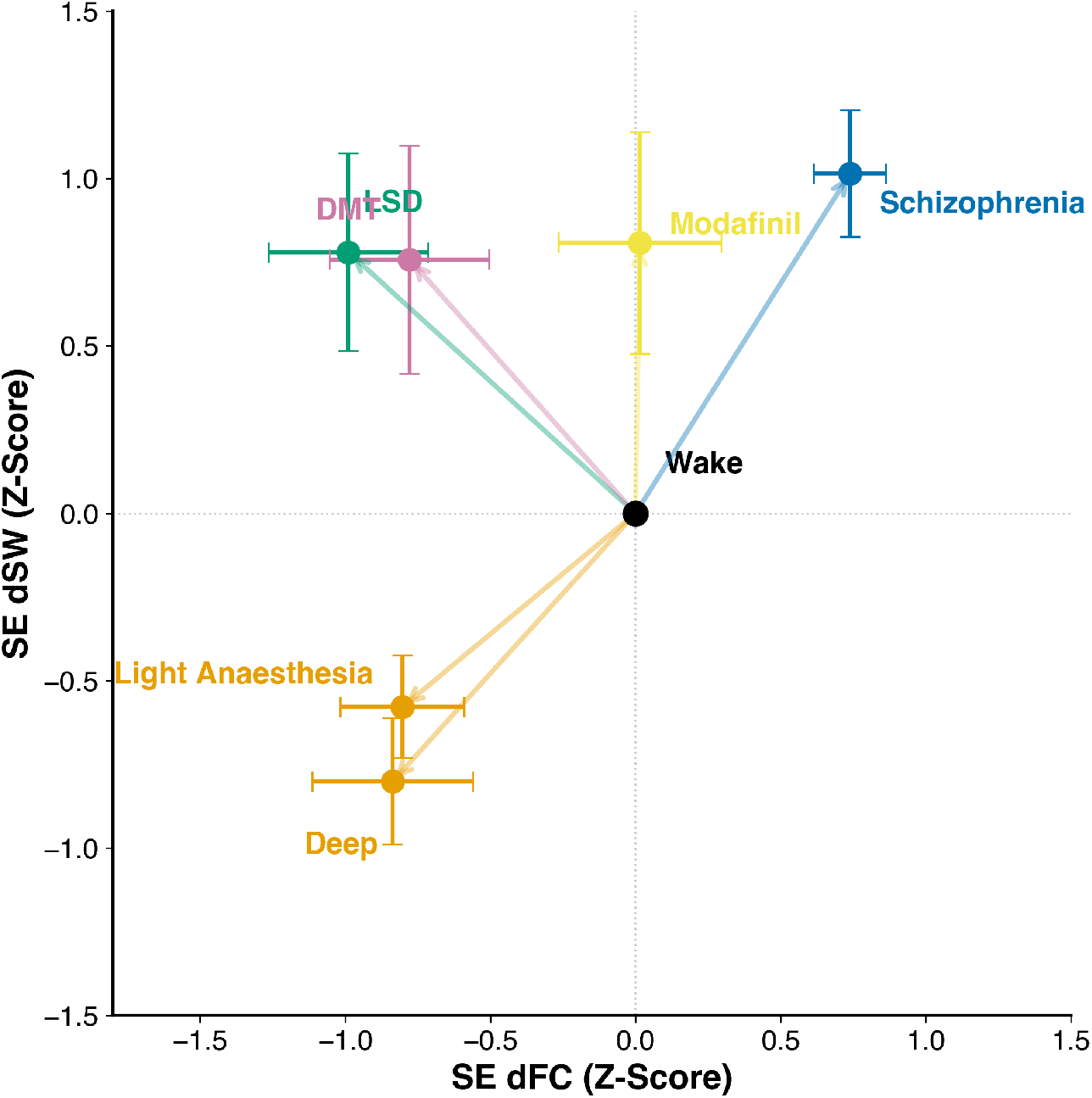
Two-dimensional state space defined by SE dSW and SE dFC. Mean SE dSW (Z-score, y-axis) is plotted against mean SE dFC (Z-score, x-axis), with both measures normalised relative to wakefulness (black point at origin). Coloured markers represent condition means; error bars indicate s.e.m.; vectors show displacement from the wakefulness baseline. Light and deep propofol anaesthesia fall in the lower-left region (reduced SE dSW and SE dFC). Schizophrenia falls in the upper-right (elevated on both axes). LSD and DMT fall in the upper-left (elevated SE dSW, reduced SE dFC), off the diagonal occupied by anaesthesia (lower-left) and schizophrenia (upper-right). Modafinil falls above the origin on SE dSW with no detectable displacement on SE dFC.

### 2.4 Network-level contributions to the entropic axis

The global entropic signature values can arise from very different patterns of network-level contribution: similar shifts in SE dSW could reflect selective changes in a few systems or distributed changes across many, and different shifts could reflect qualitatively distinct reorganisations of the same set of networks. To ask how the conditions studied here engage the seven canonical Yeo networks, we iteratively removed each network from the dynamic adjacency matrices and recomputed SE dSW on the residual system. Each network’s contribution to the global signal was estimated by the drop in SE dSW when the network was removed (NEC_k), and condition-specific effects were quantified as the change in this contribution between active state and control (ΔNEC_k; Fig. 6; see Methods). Three of the five datasets engaged higher-order association networks while sparing sensory systems. Under propofol, this pattern was present across sedation levels. Light sedation reduced ΔNEC in the Frontoparietal (p_FDR = 0.036, r = -0.76) and Salience/Ventral Attention (p_FDR = 0.039, r = -0.71) networks, with a comparable reduction in the Limbic network at trend level (p_raw = 0.029, p_FDR = 0.068, r = -0.62; Fig. 6). Deep anaesthesia engaged the same association networks more broadly —Dorsal Attention (p_raw = 0.009, r = -0.72), Frontoparietal (p_raw = 0.021, r = -0.65) and Limbic (p_raw = 0.034, r = -0.60), with a further trend in the Default Mode (p_raw = 0.065, r = -0.53) — but none of these effects survived FDR correction (all p_FDR > 0.06; Fig. 6). During post-anaesthesia recovery, the suppression of the Salience/Ventral Attention network persisted (p_FDR = 0.044, r = -0.75). Sensory networks were unaffected across all levels (Visual and Somatomotor, |r| < 0.27). Anaesthesia thus disengages association cortex rather than suppressing all systems uniformly; the effect is statistically robust at light sedation and extends, with comparable effect sizes but reduced statistical power, into deep anaesthesia. LSD and DMT produced convergent reductions in the Frontoparietal network (LSD: p_raw = 0.041, r = -0.60; DMT: p_raw = 0.049, r = -0.52), with additional reductions in the Default Mode under LSD (p_raw = 0.035, r = -0.62) and in the Limbic system under DMT (p_raw = 0.029, |r| = -0.57; Fig. 6). The psychedelic effects did not survive FDR correction, but the effect sizes are large ( r > 0.5).

**Figure 6.**
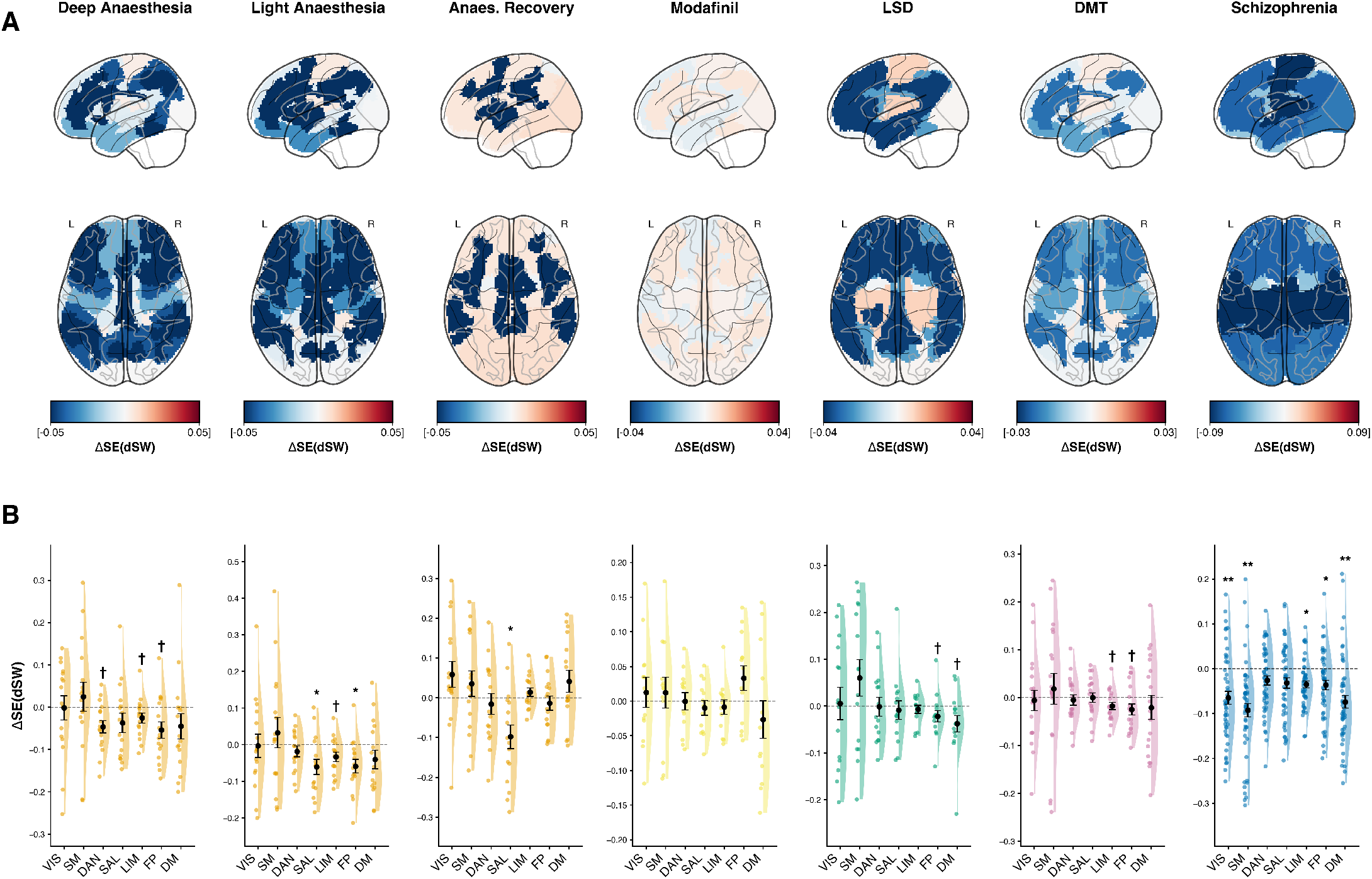
Network-level contributions to the entropic axis (leave-one-network-out analysis). Each functional network was iteratively excluded from the dynamic adjacency matrices and SE dSW was recomputed on the residual system. The Network Entropy Contribution (NEC) of each network was defined as the drop in SE dSW upon its removal; condition-specific effects are reported as ΔNEC = NEC_active - NEC_control. (A) Spatial distribution of ΔNEC. Glass-brain projections (lateral, top row; axial, bottom row, L/R as marked) show regional contributions to global SE dSW for, from left to right: deep propofol anaesthesia, light propofol anaesthesia, anaesthesia recovery, modafinil, LSD, DMT and schizophrenia. Warmer colours indicate networks whose presence increases global entropy in the active state relative to control; cooler colours indicate suppression. Note that the colour scale differs across conditions (see scale bars). (B) ΔNEC across the seven Yeo networks. Raincloud plots show ΔNEC for the Visual (VIS), Somatomotor (SM), Dorsal Attention (DAN), Salience/Ventral Attention (SAL), Limbic (LIM), Frontoparietal (FP) and Default Mode (DM) networks, with the same condition order as in (A). Black circles and error bars represent mean ± s.e.m. †p_raw < 0.05 (uncorrected), *p_FDR < 0.05, **p_FDR < 0.01.

The remaining conditions departed from this higher-order pattern in different ways. Schizophrenia produced widespread reductions in ΔNEC across five of the seven networks — Somatomotor (p_FDR = 0.003, r = -0.41), Visual (p_FDR = 0.007, r = -0.36), Default Mode (p_FDR = 0.009, r = -0.34), Limbic (p_FDR = 0.010, r = -0.32), and Frontoparietal (p_FDR = 0.042, r = -0.25) (Fig. 6). This distributed loss of per-network contribution co-occurs with the global elevation of SE dSW reported above: in schizophrenia, the elevated entropy does not arise from selective engagement of particular systems but from a generalised loss of network specificity that spares no functional division, sensory systems included. Modafinil showed no detectable network-level shifts (all p_raw > 0.22, |r| < 0.31): its modest global entropy elevation arises uniformly across systems rather than from any specific network contribution. The corresponding leave-one-network-out analysis on SE dFC (Fig. S7) showed substantially weaker network-level effects across all conditions, consistent with the global dissociation between topology and connectivity-magnitude entropy reported above: the redistribution of network contributions is a property of topological reconfiguration, not of overall coupling strength.

## 3 Discussion

The Entropic Brain Theory predicted, more than a decade ago, that levels and contents of consciousness should be ordered along a single dynamical continuum^13^. We evaluated this prediction empirically using a single dynamical index, the sample entropy of time-resolved small-world dynamics, which ordered pharmacological and clinical conditions along a continuous gradient, ranging from suppressed network reconfiguration under anaesthesia to expanded reconfiguration under psychedelics and schizophrenia. The ordering was not imposed, but emerged from a common pipeline applied to five heterogeneous datasets, and was independent of fluctuations in connectivity magnitude. Conscious states, on this evidence, are more closely tracked by the temporal flexibility of large-scale network reconfiguration than by the strength of static connectivity.

Previous work had reported results converging on this picture. Psychedelics are known to increase signal diversity^8^ and broaden connectivity repertoires^16^; anaesthesia and disorders of consciousness suppress dynamic connectivity complexity across electrophysiological and hemodynamic measures ^5,9,17,18^. Our results place these findings on a common axis, and the way they do is theoretically informative. The gradient does not index level and content as separable dimensions assigned to opposite ends; rather, it orders states by how far the temporal diversity of network con-figurations departs from ordinary wakefulness, which serves as the baseline of the axis. Both extremes are deviations from this baseline in opposite directions. At the low end, awareness is progressively suppressed and network reconfiguration becomes stereotyped and restricted, the system visiting a narrow, repetitive set of configurations. At the high end, the capacity for awareness is broadly retained while the repertoire of visited configurations expands and, in schizophrenia, becomes dysregulated. Level and content are not separated across the axis but covary along it: perturbations toward either extreme alter both, and what the axis isolates is the magnitude and direction of their joint departure from baseline.

These results provide a candidate mechanistic interpretation for the Entropic Brain Theory, and suggest why a single dynamical axis should order such heterogeneous states. Sustaining consciousness requires the brain to combine specialised local processing with efficient global communication —the integration–segregation balance that small-world organisation embodies^15^. Meeting this demand moment to moment requires the system to move through a broad but bounded repertoire of network configurations: neither so restricted that integration and segregation cannot be flexibly rebalanced, nor so unconstrained that no stable configuration is maintained. The temporal diversity of network reconfiguration is therefore not an incidental property but a precondition for the integration–segregation trade-off that conscious processing depends on. On this view a single axis is expected to order heterogeneous perturbations, because what they share is a displacement of this repertoire away from its waking regime — toward excessive order at the low end, where reconfiguration becomes stereotyped and restricted, or toward excessive disorder at the high end, where it expands and, in schizophrenia, becomes dysregulated. Anaesthesia and schizophrenia, on this reading, are opposite deviations from the same organisational principle rather than unrelated pathologies of dynamics. This framing is in turn consistent with proposals that waking consciousness operates near a critical regime^3,19^, at which the trade-off between order and disorder — and thus between integration and segregation — is most favourable, and from which states can deviate in either direction. The thalamocortical system is one plausible candidate for regulating the breadth and flexibility of cortical state dynam-ics, as the thalamus is increasingly recognised as a regulator of cortical arousal, large-scale integration, and conscious perception^20,21^. From this perspective, reductions in dynamic-network entropy under anaesthesia and elevations under psychedelics, modafinil, and schizophrenia may reflect different departures from normally bounded thalamocortical control, consistent with evidence linking thalamocortical modulation to arousal-altering and psychotomimetic states^22^. Analyses targeting thalamic and thalamocortical dynamics directly will be needed to test this mechanism.

The dissociation between sample entropy and functional connectivity variability under psychedelics is particularly informative. Psychedelics increased the temporal diversity of large-scale network topology while suppressing the variability of overall coupling strength: the same compound that expanded the mesoscopic repertoire drove global connectivity magnitude in the opposite direction. This opposing modulation distinguishes the psychedelic state from other perturbations along the axis. Anaesthesia suppresses both broadly; schizophrenia amplifies both broadly; psychedelics split the two, expanding topological diversity while suppressing connectivity-magnitude variability. The picture is consistent with evidence that psychedelics expand the brain’s dynamical repertoire toward criticality^23^ and flatten the energy landscape of state transitions^24^, rather than producing a uniform increase in dynamical activity. Such dissociations between complexity and entropy are not unique to psychedelics: in a voluntarily induced non-ordinary state characterised by increased visual content, statistical complexity increased while permutation entropy decreased rela-tive to baseline^25^— moving in the opposite direction to the psychedelic signature despite both states sharing expanded experiential content. Together these results indicate that conscious states are not characterised by a single dynamical quantity but by specific configurations across complementary descriptors, and that the position of a state along the entropic axis does not predict its full dynamical profile.

A further implication follows from where the conditions fall along the axis. The typical conscious state appears to inhabit a narrow regime of dynamical entropy. Too little SE yields the stereotyped, unresponsive dynamics of anaesthesia; too much yields the dysregulated dynamics observed in schizophrenia. Psychedelics push brain dynamics beyond the typical waking range while preserving insight into the altered nature of the state, distinguishing the psychedelic condition from psychosis even though both expand the entropic repertoire. Healthy waking consciousness sits between these extremes, in a regime flexible enough to support increases in the richness of contents experienced but constrained enough to preserve coherence — consistent with proposals positing that the conscious brain operates near a critical regime^26^, where the trade-off between order and disorder is most favourable.

Several limitations qualify these results. First, all data were acquired at rest, which is suboptimal for probing the qualitative organisation of conscious experience: links between the entropic axis and content reported here are indirect, inferred from differences across conditions rather than from variation in measured content. Future work could test whether SE dSW predicts more direct mark-ers of content — inter-subject synchrony during narrative comprehension^27^ or experience-sampling probes during waking cognition. Second, the present design lacks a clinical population with graded reductions in awareness. Disorders of consciousness offer the most direct behavioural anchor for the low-entropy end of the axis^28^ and would allow the gradient to be validated against measures such as the Coma Recovery Scale–Revised. Third, the five datasets differ in scanner, site, and protocol. Most of these differences did not affect the gradient: the healthy controls of the schizophrenia cohort, the LSD and DMT placebos, and the propofol wake-baseline cohort showed closely overlapping SE dSW values (Fig. S4). The modafinil cohort was the exception, showing a uniform acquisition-related offset (Fig. S4); because the modafinil and placebo groups were acquired within the same cohort, this offset is shared across the contrast and does not affect the modafinil–placebo comparison. Largescale validation across coordinated multi-site cohorts with harmonised acquisition parameters would establish the robustness of the axis. Fourth, our analyses are observational. Although the within-subject reversibility of the anaesthesia effect — entropy falling under propofol and returning to baseline upon recovery — indicates that SE dSW tracks the active pharmacological state rather than a stable trait or acquisition artefact, perturbational approaches such as TMS-EEG indices of complexity^5,6^ or dose-response pharmaco-logical designs will be needed to establish whether changes in topological flexibility are causal with respect to changes in conscious states. These limitations notwithstanding, the projection of pharmacologically and clinically heterogeneous conditions onto a single topological-entropy axis provides the cross-condition test that the Entropic Brain Theory has lacked. That anaesthesia, psychedelics, modafinil and schizophrenia order along one dimension indicates that the temporal diversity of large-scale network reconfiguration captures a genuine axis of conscious-state variation, and offers a tractable target for the perturbational work needed to establish its causal status

## 4 Methods

### 4.1 Datasets and experimental conditions

Five fMRI resting-state datasets, spanning four pharmacological and one clinical perturbation, were analysed with an identical pipeline. All original studies obtained ethical approval from their respective committees and written informed consent from participants. Table 1 summarises acquisition parameters.

**Table 1.** Acquisition parameters and sample sizes across the five datasets. Each row lists a dataset analysed with the common analytical pipeline. N indicates the number of participants entering the analysis after the exclusions reported in the original studies; for the schizophrenia dataset, N is the combined patient-plus-control sample (50 patients, 50 controls). Age range is given as the reported range or as mean ± s.d., depending on the original study. TR is in seconds and scan duration in minutes; for the propofol dataset, the duration corresponds to the deep-anaesthesia condition. The reference column cites the original publication of each dataset. Full descriptions are given in Methods.

| Dataset | N | Age (range or mean ± s.d.) | Scanner | TR (s) | Scan duration (min) | Reference* |
| --- | --- | --- | --- | --- | --- | --- |
| LSD | 15 | 30.5 ± 8.0 | 3 T GE HDx | 2.0 | 14.0 | <sup>3</sup> (Carhart-Harris 2016 PNAS) |
| DMT | 14 | 33.5 ± 7.9 | 3 T Siemens Verio | 2.0 | 28.0 | <sup>31</sup> (Timmermann 2023 PNAS) |
| Modafinil | 24 | 25-35 | 3 T Philips Achieva | 1.67 | 12.1 | <sup>22</sup> (Delli Pizzi 2025 NeuroImage) |
| Propofol | 17 | 18-40 | 3 T Siemens | 2.0 | 8.5 | <sup>30</sup> (Naci 2018 Sci Rep) |
| Schizophrenia | 100 | 34.4 ± 9.2 | 3 T Siemens Trio | 2.0 | 5.1 | <sup>31</sup> (Poldrack 2016) |

#### 4.1.1 LSD

A double-blind, placebo-controlled, within-subject study at Imperial College London ^3^. Fifteen healthy volunteers completed two scanning sessions (LSD and placebo) separated by at least two weeks, in balanced order. LSD (75 *µ*g in 10 mL saline) or placebo (10 mL saline) was administered intravenously over 2 minutes. Resting-state BOLD fMRI was acquired in two runs totalling 14 minutes, beginning 135 minutes after administration, with eyes closed and no explicit task. Imaging parameters acquisition: 3 T GE HDx, T2-weighted gradient-echo EPI, TR = 2000 ms, TE = 35 ms, voxel size 3.4 × 3.4 × 3.4 mm^3^, 35 slices, flip angle 90°, FOV 220 mm, matrix 64 *×* 64.

#### 4.1.2 DMT

A single-blind, placebo-controlled, counterbalanced study at Imperial College London^31^, approved by the UK National Research Ethics Committee London–Brent and the Health Research Authority and conducted under a Home Office license for Schedule 1 substances. Of the 20 participants originally enrolled, 14 were retained for analysis following the motion exclusion criteria of the original paper^31^. Participants underwent two sessions (DMT and placebo) separated by two weeks. On each session, 20 mg of intravenous DMT fumarate (in 10 mL sterile saline) or placebo (saline only) was administered over 30 s during a 28-min resting-state fMRI acquisition. Scans were obtained with eyes closed (eye mask), with concurrent EEG. Acquisition: 3 T Siemens Magnetom Verio, T2-weighted EPI, TR = 2000 ms, TE = 30 ms, voxel size 3 × 3 × 3 mm^3^, 35 slices.

#### 4.1.3 Modafinil

A double-blind, placebo-controlled study^22^. 24 healthy volunteers were randomly assigned to receive either 100 mg oral modafinil (N=13) or placebo (N=11). Resting-state scans were acquired 2 hours post ingestion with eyes open, fixating on a central point. Acquisition: 3 T Philips Achieva, T2-weighted gradient-echo EPI, TR = 1670 ms, TE = 30 ms, voxel size 4 *×* 4 *×* 4 mm^3^, 30 axial slices, flip angle 75°, FOV 256 mm, matrix 64 *×* 64; three runs of 145 volumes per participant.

#### 4.1.4 Propofol anaesthesia

The anaesthesia data was collected at Western University^30^. Seventeen healthy adults were scanned across four conditions: awake baseline, light propofol anaesthesia, deep propofol anaesthesia, and post-anaesthesia recovery. Propofol was administered intravenously following Canadian Anesthesiology Society guidelines, with continuous monitoring of blood pressure, heart rate, oxygen saturation, and end-tidal CO_2_. Propofol infusion commenced at an effect-site concentration of 0.6 *µ*g/mL and was titrated upward until the target level of deep sedation was achieved and maintained during scanning. Acquisition: 3 T Siemens, TR = 2000 ms, TE = 30 ms, 3 mm^3^ isotropic voxels, 256 volumes per run.

#### 4.1.5 Schizophrenia

The schizophrenia data was collected at UCLA^31^. Fifty patients with schizophrenia and 50 matched healthy controls were included. Ethical approval was granted by the UCLA Institutional Review Board. Resting-state fMRI was acquired with eyes open for 304 s. Acquisition: 3 T Siemens Trio (syngo MR B15/B17), T2-weighted EPI, TR = 2000 ms, TE = 30 ms, voxel size 3 *×* 3 *×* 4 mm^3^, 34 slices, flip angle 90°.

### 4.2 Preprocessing

The preprocessing pipeline was performed using the default pipeline in CONN toolbox^32^ version 22.a. Functional images were realigned to the first scan of each session and unwarped to account for magnetic field inhomogeneities, followed by slice-timing correction (centered at the middle of the TR). Potential outliers were identified using the Artifact Detection Tool (ART) with a framewise displacement (FD) threshold of 0.4 mm. Structural and functional data were normalized to MNI152 space through a unified segmentation and normalization procedure, resampled to 2 mm isotropic voxels, and spatially smoothed using a 6 mm FWHM Gaussian kernel. To minimize non-neural noise, the denoising step included the aCompCor strategy, regressing out five principal components each from the white matter and cerebrospinal fluid masks. Additionally, 12 motion parameters (6 translation/rotation parameters plus their first-order temporal derivatives) and the identified outlier volumes (scrubbing) were regressed out. The BOLD time series were subjected to linear detrending, and band-pass filtering [0.008–0.09 Hz] to isolate low-frequency fluctuations. The only exception was the LSD dataset, for which we utilized the preprocessed and denoised BOLD time series provided by the original authors, as the raw data were not available.

To ensure that head motion did not confound condition contrasts, mean framewise displacement (FD_mean) was extracted using MRIQC. We evaluated the effect of head motion on sample entropy using linear models controlling for condition. Data for the LSD dataset was not included in this analysis as it was obtained already preprocessed. In the anaesthesia, DMT, and modafinil datasets, the effect of head motion on signal complexity was not statistically significant (all p > 0.14). In the schizophrenia dataset, head motion showed a weak negative association with sample en-tropy (raw p = 0.035); crucially, however, the main effect of clinical diagnosis remained highly significant (p < 0.001) even after regressing out head motion. These findings confirm that head motion is not the primary driver of the observed topological complexity shifts (Table S1).

### 4.3 Parcellation

For each subject, BOLD time series were extracted from regions of interest defined by AAL90. Robustness to parcellation choice was assessed by repeating the entire pipeline using a combination of Melbourne subcortical^33^ and the Schaefer cortical^34^ parcellations, similarly to previous studies^15,35^. The leave-one-network-out analysis (see below) used the Schaefer cortical parcellation with parcels assigned to the seven canonical Yeo resting-state networks^36^.

### 4.4 Dynamic functional connectivity

We followed the analytical framework by Coppola et al.^15^ to characterise time-resolved changes in large-scale network organisation. For each participant and condition, sliding windows of approximately 48 s were applied to the preprocessed BOLD time series: 24 TRs for datasets with TR = 2 s (LSD, DMT, propofol, schizophrenia) and 28 TRs for the modafinil dataset (TR = 1.67 s). Windows were advanced in steps of one TR, yielding partially overlapping segments covering the entire scan. Each window was tapered with a Hamming window to down-weight boundary time points^37^. Within each window, Pearson correlation coefficients between all pairs of ROIs produced a symmetric functional connectivity (FC) matrix at each time point. The diagonal was set to zero and only positive correlations were retained. This choice reflects two considerations: the graph-theoretical metrics used here (clustering coefficient, characteristic path length, small-worldness) are defined for positively-weighted graphs; and the biological interpretation of negative BOLD correlations remains ambiguous, as they can arise either from genuine anticorrelation or from preprocessing-related effects^38^.

In parallel, a scalar index of dynamic FC (dFC) was derived per window by averaging the positive entries of the unthresholded FC matrix. The resulting dFC time series captures moment-to-moment changes in the overall strength of functional coupling^15,39^.

### 4.5 Graph construction and small-world topology

Graph-theoretical analyses were carried out on weighted, undirected networks derived from the windowed FC matrices, with ROIs treated as nodes and correlation coefficients as edge weights^38^. Because no single thresholding strategy is universally optimal^35,38^, we adopted a proportional thresholding scheme and explicitly tested robustness across a range of densities. For each FC matrix, edges were ranked by weight, and the strongest connections were retained up to five target densities sampled from 0.05 to 0.25 in steps of 0.05 (i.e., the top 5–25% of edges). This generated five thresholded graphs per window and participant.

This multi-density approach mitigates dependence on any single threshold^15,35^. Two further features of the analysis address concerns with proportional thresholding in clinical comparisons^39,40^: first, edge weights were retained rather than binarised, reducing the contribution of weaker edges to graph metrics; and second, the entropy of mean dFC was independently quantified (see below), allowing topological effects to be dissociated from differences in FC amplitude. For each thresholded matrix, we computed the clustering coefficient (the tendency of a node’s neighbours to be interconnected) and the characteristic path length (the mean shortest-path distance between all node pairs)^38,41^. Both were normalised against appropriate reference networks. We then computed a small-world propensity index that compares the clustering coefficient to a lattice graph and the characteristic path length to a degree-matched random graph^41,42^. This index lies between 0 and 1 and quantifies how closely a given network approximates the small-world regime of high clustering and short paths.

Graph computations were implemented in MATLAB and Python using the Brain Connectivity Toolbox^38^ and custom scripts.

### 4.6 Sample entropy

Sample entropy (SE) was applied to the time series of dynamic small-worldness (dSW) and to the dFC time series, following the implementation of Coppola et al. ^15^. SE quantifies how often short temporal patterns that are similar within a tolerance remain similar when extended by one time point, excluding trivial self-matches^43,44^. Higher SE values indicate less recurrence of temporal motifs and greater temporal diversity; lower values indicate more stereotyped dynamics.

Parameters were set following standard practice in fMRI graph dynamics^15^: embedding dimension *m* = 2; tolerance *r* = 0.2 *×* within-series standard deviation; natural logarithms. Scaling the tolerance by the within-series SD makes the criterion invariant to absolute signal amplitude.

For each subject, parcellation, and density, SE was computed on the dSW time course. SE values were then averaged across the five densities to obtain a single index of dynamic-network entropy (SE dSW) per subject and condition, reducing sensitivity to any single edge density. The same procedure was applied to the dFC time series, yielding SE dFC as a complementary index of the irregularity of overall coupling magnitude.

### 4.7 Leave-one-network-out (LONO) decomposition

To identify which functional systems contribute to condition-related shifts in SE dSW, we extended the analysis with a leave-one-network-out decomposition on the Schaefer cortical parcellation, with parcels assigned to the seven canonical Yeo resting-state networks. For each network k, all parcels belonging to that network were iteratively excluded from the dynamic adjacency matrices across all sliding windows, yielding a reduced six-network subgraph at each time point. The full pipeline (proportional thresholding across five densities, computation of dSW, SE estimation) was then re-applied to the residual system. The Network Entropy Contribution (NEC) of each network was defined as:

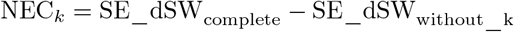

Positive values indicate that the presence of network k increases whole-brain entropy. Condition-related changes were quantified as:

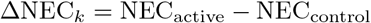

### 4.8 Statistical analysis

Effects of condition on SE dSW and SE dFC were tested using linear mixed-effects models with random intercepts for each subject to account for repeated measures in within-subject designs (psychedelic and propofol datasets). For independent between-subject designs (schizophrenia and modafinil datasets), standard ordinary least squares (OLS) regression was used. Fixed effects were defined by condition in all models. Models were fitted by restricted maximum likelihood (REML) using the statsmodels (0.14.4) Python (3.12) package. For each contrast we report the regression coefficient (*β* for single-dataset models, Coef for the unified cross-dataset model), 95% confidence interval, and two-tailed p-value.

For the unified gradient analysis across datasets (Fig. 3), SE dSW was first standardised within each study to place all datasets on a common scale, and each perturbation was then expressed as its standardised change relative to its within-study baseline (ΔSE dSW; active minus control for the pharmacological and clinical datasets, and each sedation level minus the awake baseline for the propofol dataset), removing site- and acquisition-related differences in baseline across studies. These standardised differences were entered into a single mixed-effects model with all conditions as categorical fixed effects and no global intercept (Delta_Z 0 + C(Condition) + (1 | Subject)), with subject-level random intercepts retaining the within-subject structure of each original study. This specification estimates the standardised deviation of each state from baseline wakefulness independently. The corresponding raw, un-differenced SE dSW values are reported in Fig. S4 and yield a qualitatively identical ordering.

For the LONO analysis (ΔNEC across the seven Yeo networks), differences from zero were tested using two-tailed Wilcoxon signed-rank tests for within-subject comparisons (psychedelics, and propofol datasets) and Mann-Whitney U tests for the between-subject comparison (schizophrenia and modafinil). Effect sizes were quantified by rank-biserial correlation (r). The seven per-condition tests were corrected for multiple comparisons using the Benjamini–Hochberg false discovery rate procedure (*α* = 0.05). We report both FDR-corrected (p_FDR) and uncorrected (p_raw) values to allow readers to assess the convergence of effects that did not survive correction.

## Code and data availability

The datasets analyzed in this study are available from public repositories or upon request from the authors. The schizophrenia dataset is available on OpenNeuro under the accession number ds000030 (https://openneuro.org/datasets/ds000030). The LSD dataset is available on OpenNeuro under the accession number ds003059 (https://openneuro.org/datasets/ds003059). The Modafinil and DMT datasets are not publicly available and were obtained directly from the respective authors; requests for access to these datasets should be directed to them. Analysis code is available at https://github.com/GalvanRial-DS/entropic-axis.

## Acknowledgments

This research was supported by Agencia Nacional de Promoción Científica y Tecnológica, Argentina (Grants #2018-03614, CAT-I-00083) and Stic Amsud project (CONN-COMA, 2023). PB and TMO were supported by the National Scientific and Technical Research Council (CONICET - Argentina). D.S.G.R. was supported by a doctoral fellowship from CONICET, Argentina.

## Statement of contributions (CRediT)

Conceptualization was carried out by P.B. and E.A.S. Methodology was designed by P.B., E.A.S. and D.S.G.R. Software was developed by D.S.G.R. and G.A.D.B. Validation was performed by D.S.G.R. Formal analysis was carried out by D.S.G.R. and T.M.O. Investigation was conducted by D.S.G.R. Resources were provided by L.N., S.S., C.T., R.C.-H. and E.A.S. Visualization was produced by D.S.G.R. The original draft was written by D.S.G.R. and P.B. All authors reviewed and edited the manuscript. Supervision was provided by P.B. Project administration was carried out by P.B. Funding was acquired by P.B. and E.A.S.

## Supplementary Material

**Table S1.** Head-motion control. Effect of mean framewise displacement (FD_mean) on sample entropy, estimated separately for each dataset after controlling for condition. For within-subject datasets a linear mixed-effects model with subject-level random intercepts was used (MixedLM); for between-subject datasets an ordinary least-squares model was used (OLS). N is the number of observations entering each model. For each model we report the regression coefficient for FD_mean (*β*), its standard error (SE) and the uncorrected two-tailed p-value. In most datasets, head motion did not significantly predict signal complexity. While head motion had a statistically significant effect in the schizophrenia dataset, the main effect of condition remained highly robust (p < 0.001) despite covariate control. (Note: The LSD dataset was excluded from this analysis because only motion-corrected BOLD time series were available from the original authors)

| Dataset | Model | N | $\beta$ | SE | p |
| --- | --- | --- | --- | --- | --- |
| Anaesthesia | MixedLM | 68 | -0.1081 | 0.063 | 0.4579 |
| Modafinil | OLS | 24 | 0.4620 | 0.3058 | 0.1457 |
| DMT | MixedLM | 28 | -0.0243 | 0.3456 | 0.9439 |
| Schizophrenia | OLS | 100 | -0.2203 | 0.1033 | 0.0354 |

**Figure S1.**
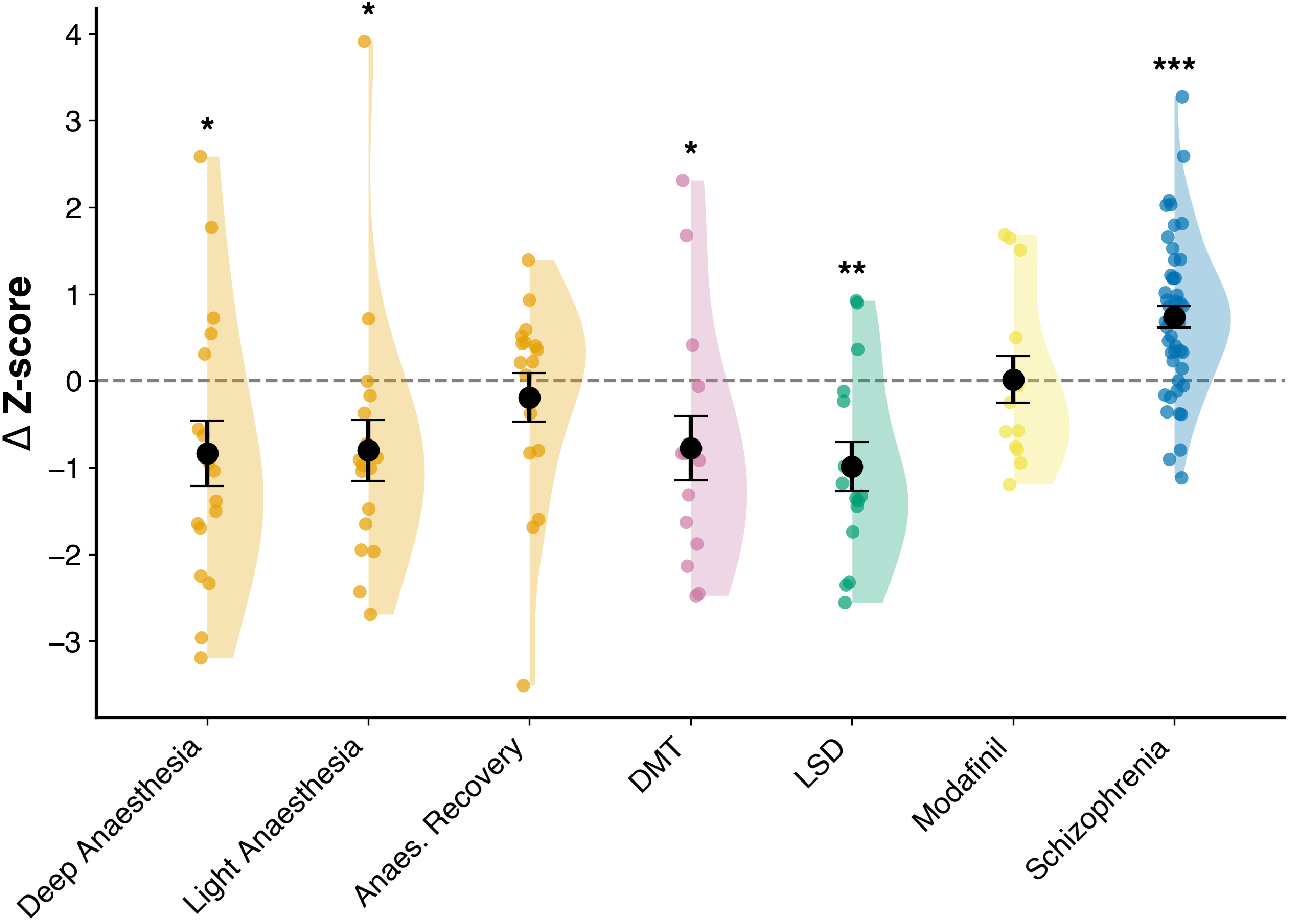
SE dFC across perturbations of consciousness. Same display and condition order as Fig. 4, but showing the standardised change in SE dFC (Δ Z-score) — the sample entropy of the dynamic functional connectivity time series, which indexes the temporal irregularity of overall coupling magnitude — instead of SE dSW. Schizophrenia anchors the high end of the axis, as for SE dSW; at the opposite extreme, however, the psychedelic conditions join the anaesthesia conditions at the low end (DMT lowest, below anaesthesia), rather than occupying the high end they reach for SE dSW. This is the topology–connectivity dissociation reported in the main text. Individual dots represent single-subject data; error bars indicate s.e.m. †p < 0.10, *p < 0.05, **p < 0.01, ***p < 0.001.

**Figure S2.**
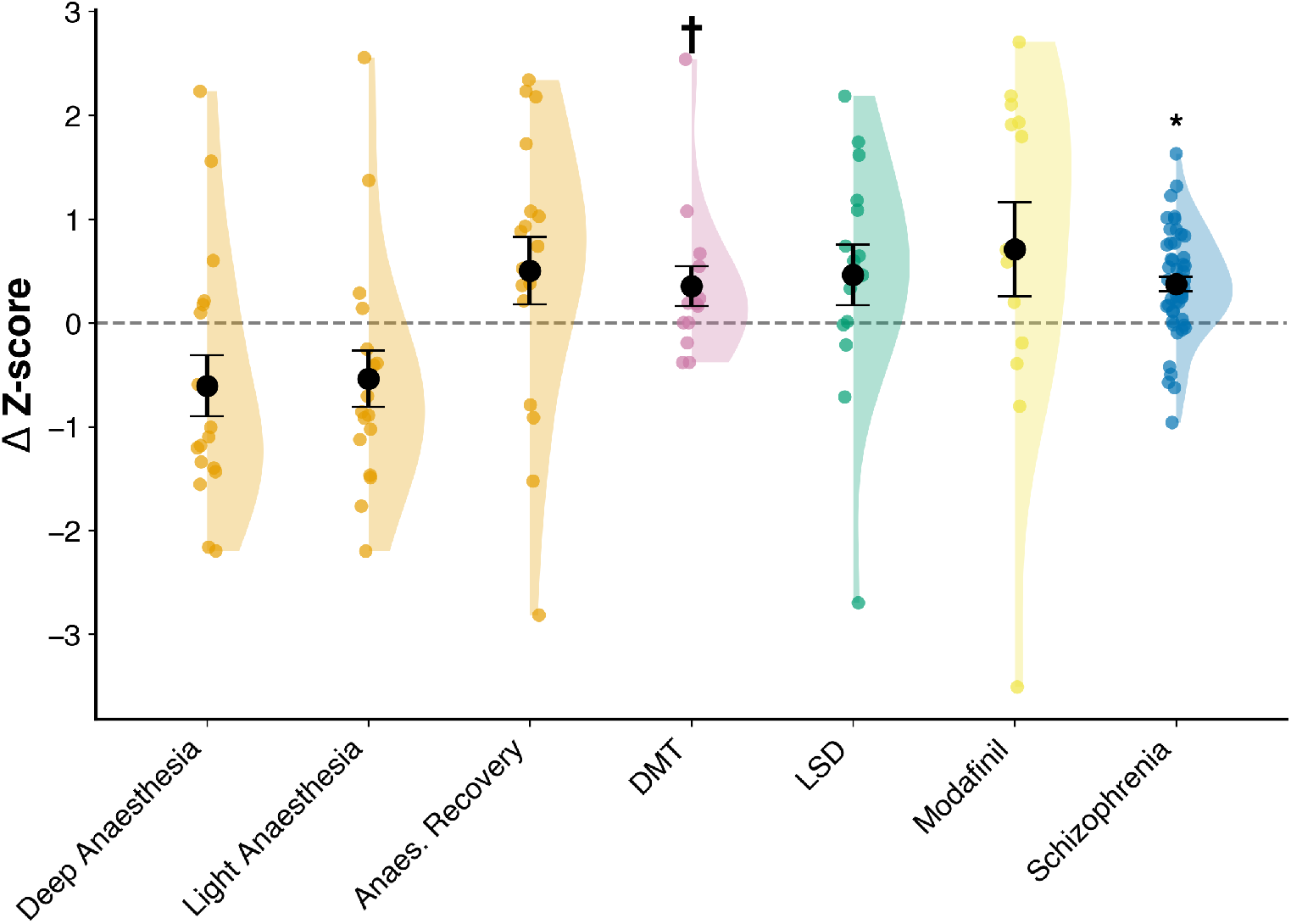
Entropic gradient (SE dSW) computed with the Schaefer cortical and Melbourne subcortical parcellation. Replication of Fig. 4 with an alternative parcellation. For each perturbation, SE dSW was standardised within study and its change relative to the within-study baseline is shown (ΔSE dSW; active minus control, or each sedation level minus awake baseline), in standardised units (Δ Z-score). Conditions are shown in the same order as in Fig. 4. The dashed line marks no change from baseline (Δ = 0). The ordering is qualitatively preserved relative to the AAL parcellation: anaesthesia conditions fall below baseline, anaesthesia recovery returns toward baseline, and DMT, LSD, modafinil and schizophrenia fall above it. Individual dots represent single subjects or sessions; black circles denote group means; error bars indicate s.e.m. †p < 0.10, *p < 0.05, **p < 0.01, ***p < 0.001.

**Figure S3.**
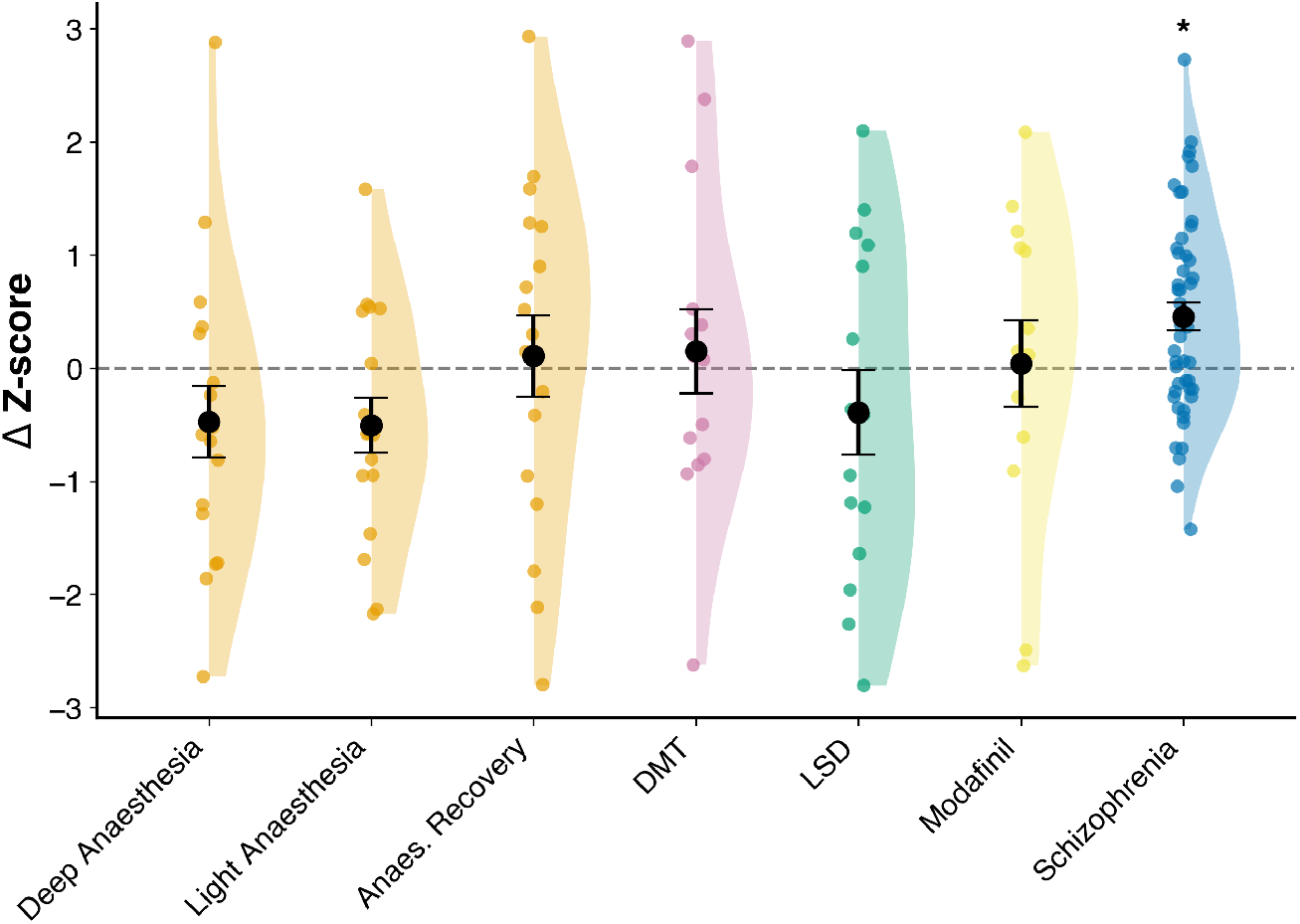
SE dFC gradient computed with the Schaefer cortical and Melbourne subcortical parcellation. Replication of Supplementary Fig. 1 with an alternative parcellation. For each perturbation, the standardised change in SE dFC relative to the within-study baseline is shown (Δ Z-score), with conditions in the same order as in Fig. 4. The dashed line marks no change from baseline (Δ = 0). As with the AAL parcellation, schizophrenia falls at the high end of the SE dFC axis while the psychedelic conditions sit at the low end, opposite to their position on the SE dSW gradient — the topology–connectivity dissociation reproduced under an alternative parcellation. Individual dots represent single subjects or sessions; black circles denote group means; error bars indicate s.e.m. †p < 0.10, *p < 0.05, **p < 0.01, ***p < 0.001.

**Figure S4.**
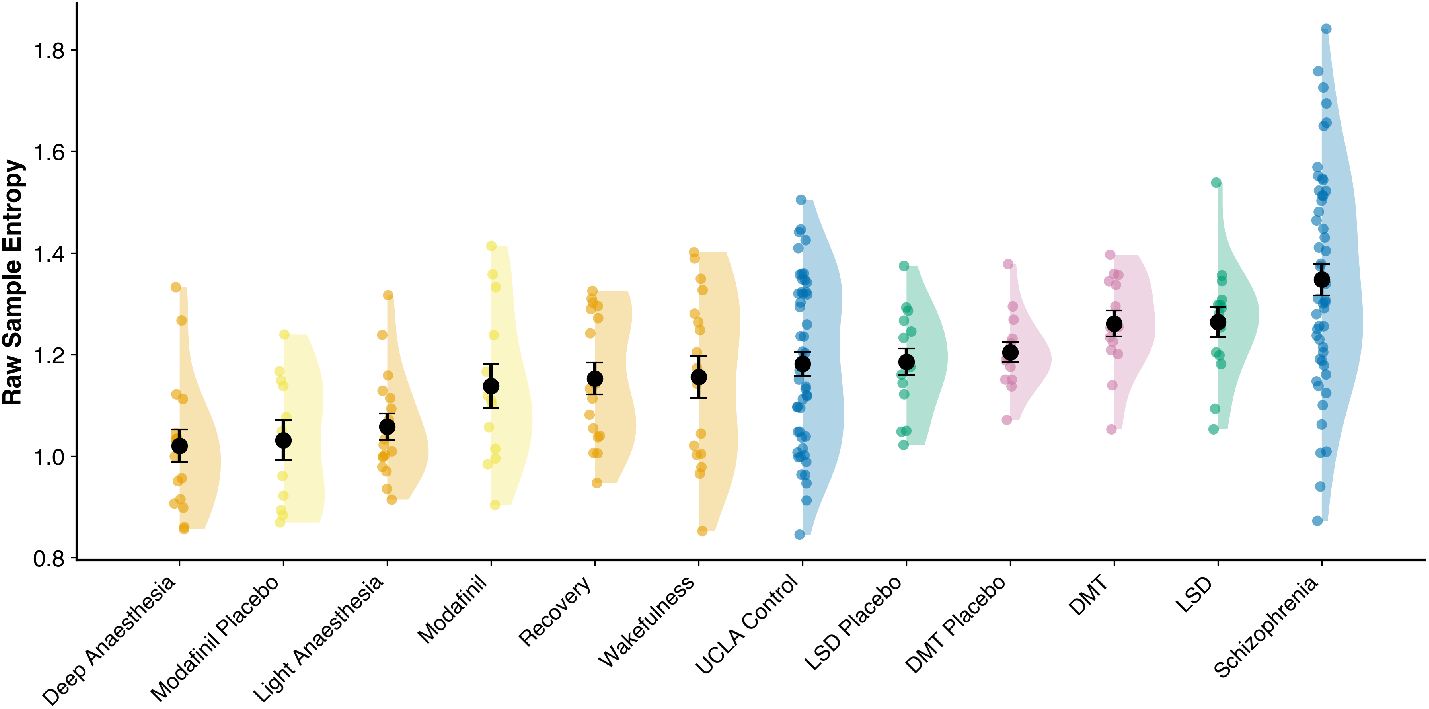
Raw SE dSW across all conditions and baselines. Distribution of SE dSW for every condition and baseline analysed, shown as raw (non-standardised, non-differenced) values, ordered by group mean from lowest to highest. Unlike Fig. 4, each placebo, control and baseline group is plotted as a separate distribution rather than absorbed into a within-study difference, so that absolute SE dSW values can be compared directly across studies. The healthy resting-state conditions acquired at different sites — wakefulness (propofol baseline), anaesthesia recovery, schizophrenia controls, and the LSD and DMT placebos — converge on closely overlapping SE dSW values, whereas the modafinil placebo sits below them, reflecting a uniform acquisition-related offset in that cohort that does not affect within-study contrasts. The overall ordering reproduces the standardised gradient of Fig. 4: deep anaesthesia lowest, schizophrenia highest, with DMT and LSD near the upper end. Individual dots represent single subjects or sessions; black circles denote group means; error bars indicate s.e.m.

**Figure S5.**
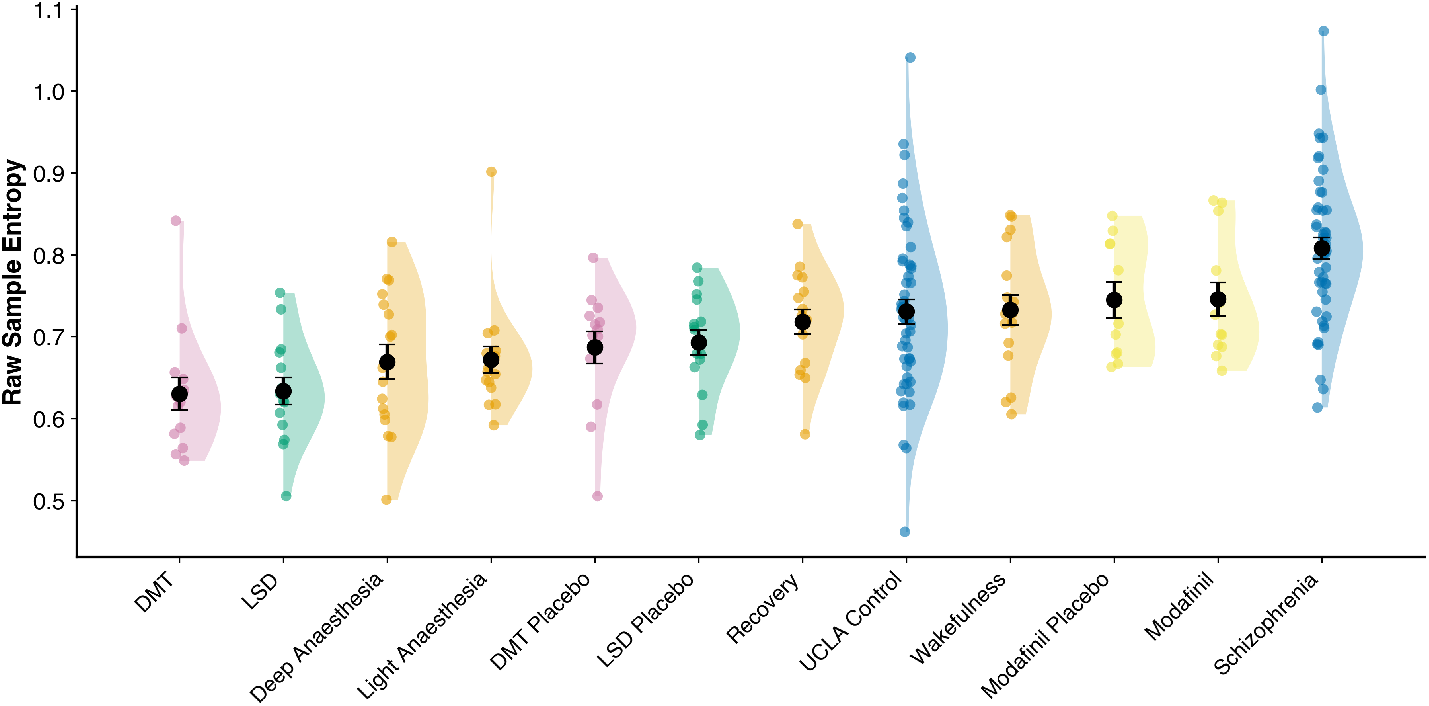
Raw SE dFC across all conditions and baselines. Distribution of SE dFC for every condition and baseline analysed, shown as raw (non-standardised, non-differenced) values, ordered by group mean from lowest to highest. As in Supplementary Fig. 4, each placebo, control and baseline group is plotted separately so that absolute SE dFC values can be compared directly across studies. Schizophrenia occupies the high end, as for SE dSW; at the opposite extreme, however, DMT and LSD fall at the low end of the SE dFC axis — below their own placebos and below the healthy resting-state baselines — the reverse of their position on the raw SE dSW gradient (Supplementary Fig. 4). This is the topology–connectivity dissociation of the main text, shown here on raw values. Individual dots represent single subjects or sessions; black circles denote group means; error bars indicate s.e.m.

**Figure S6.**
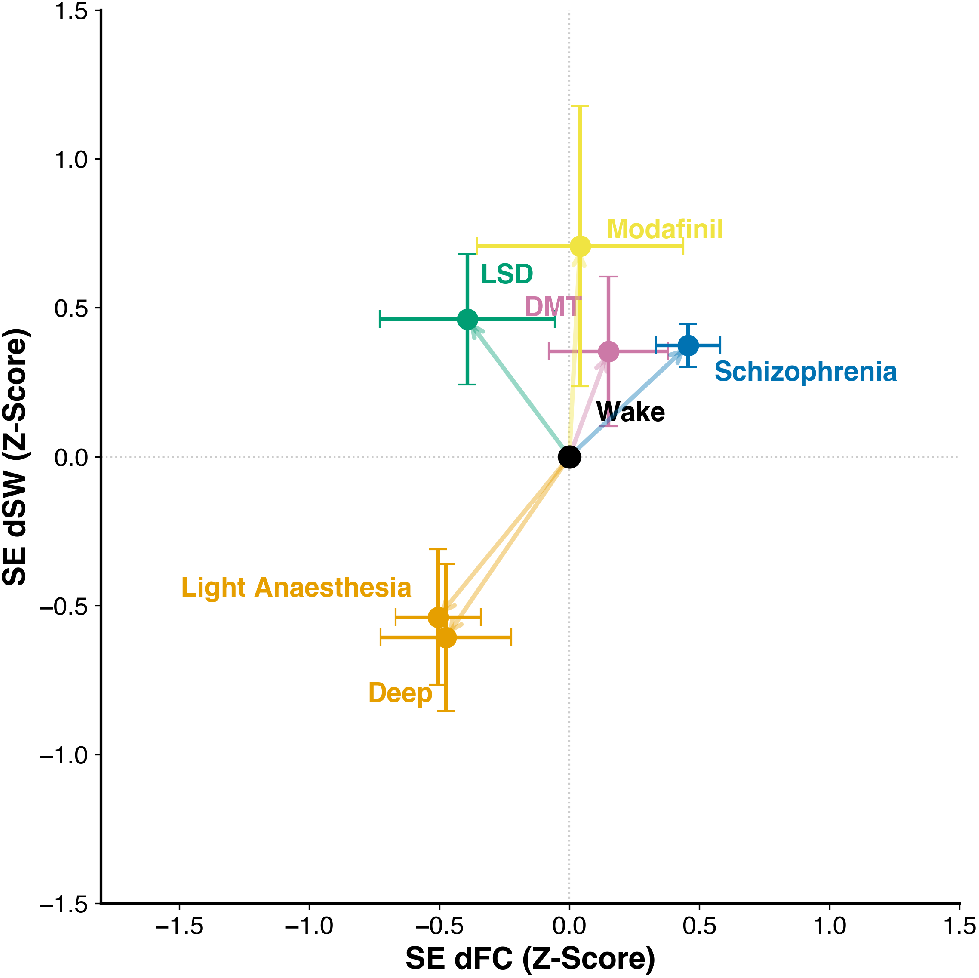
Two-dimensional state space defined by SE dSW and SE dFC, computed with the Schaefer cortical and Melbourne subcortical parcellation. Replication of Fig. 5 with an alternative parcellation. SE dSW (Z-score) is plotted on the y-axis and SE dFC (Z-score) on the x-axis; both measures are normalised relative to wakefulness (black marker at origin). Markers indicate condition means, error bars indicate s.e.m., and vectors show displacement from wakefulness. The overall arrangement is largely preserved: anaesthesia falls in the lower-left, schizophrenia in the upper-right, and modafinil is elevated on SE dSW alone. The psychedelic conditions remain elevated on SE dSW; their displacement on SE dFC is less pronounced than with the AAL parcellation — clearest for LSD, whereas DMT shows little net change on the SE dFC axis. The qualitative separation of conditions along SE dSW is thus robust to parcellation choice, while the topology–connectivity dissociation is reproduced more strongly for LSD than for DMT.

**Figure S7.**
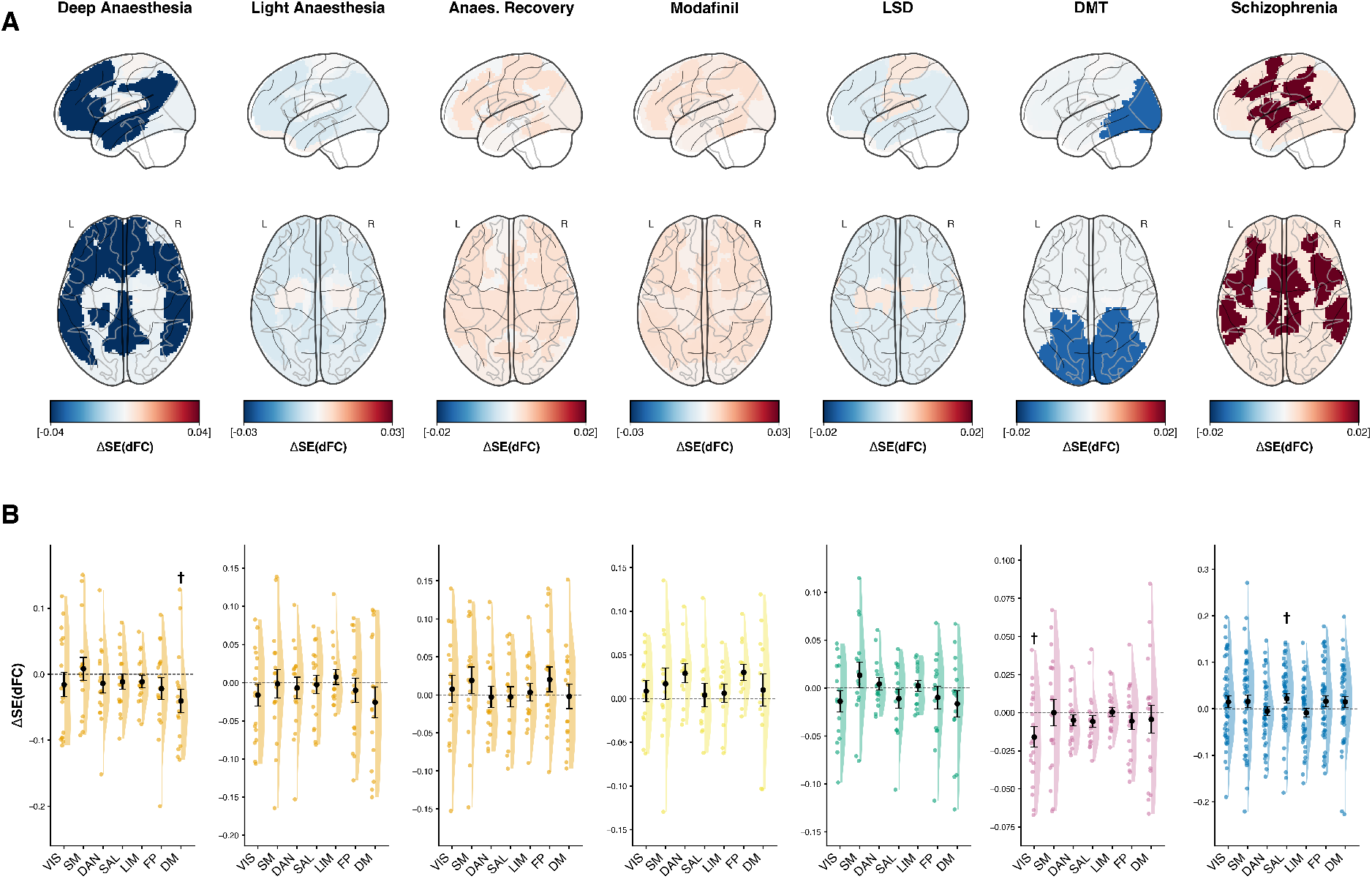
Network-level contributions to SE dFC (leave-one-network-out analysis). Same analysis as Fig. 6 applied to SE dFC instead of SE dSW. Each functional network was iteratively excluded from the dynamic adjacency matrices, and SE dFC was recomputed on the residual system. The Network Entropy Contribution (NEC) of each network is the drop in SE dFC upon its removal; condition-specific effects are reported as ΔNEC = NEC_active - NEC_control. (A) Spatial distribution of ΔNEC. Glass-brain projections (lateral, top row; axial, bottom row, L/R as marked), from left to right: deep propofol anaesthesia, light propofol anaesthesia, anaesthesia recovery, modafinil, LSD, DMT and schizophrenia. Warmer colours indicate networks whose presence increases global SE dFC in the active state relative to control; cooler colours indicate suppression; the colour scale differs across conditions (see scale bars). (B) ΔNEC across the seven Yeo networks: Visual (VIS), Somatomotor (SM), Dorsal Attention (DAN), Salience/Ventral Attention (SAL), Limbic (LIM), Frontoparietal (FP) and Default Mode (DM), with the same condition order as in (A). Black circles and error bars represent mean ± s.e.m. †p_raw < 0.05 (uncorrected), *p_FDR < 0.05, **p_FDR < 0.01. Network-level effects are substantially weaker than for SE dSW (Fig. 6) across all conditions, consistent with the global topology–connectivity dissociation reported in the main text.

